# Renal nutrient loss promotes metabolic health at the expense of increased vulnerability to acute kidney injury

**DOI:** 10.64898/2026.09.24.754002

**Authors:** Harry B. Cutler, Alexis Diaz Vegas, Oliver K. Fuller, Kristen C. Cooke, Stewart W.C. Masson, Søren Madsen, Jacqueline Stöckli, Sanae Ashitani, Kaori Igarashi, Tomoyoshi Soga, Liang Gao, Federico Torta, David E. James

## Abstract

Obesity is a major risk factor for interrelated cardiorenal metabolic diseases including type 2 diabetes, heart failure and chronic kidney disease^1–4^. Although obesity develops when energy intake chronically exceeds energy expenditure, individuals vary markedly in the amount of adiposity and metabolic dysfunction that develop in response to comparable caloric excess. Controlled overfeeding studies demonstrate several-fold variation in weight gain between individuals^5^, while susceptibility to metabolic dysfunction also differs substantially between populations^6^, suggesting considerable variation in how excess nutrients are partitioned. Energy balance is conventionally considered in terms of intake and expenditure, yet energy-containing nutrients can also be eliminated from the body. The kidney is uniquely positioned to regulate this third component of energy balance by determining whether filtered metabolites are reabsorbed or excreted. The metabolic benefits of SGLT2 inhibition demonstrate that reducing renal nutrient reabsorption can favourably alter systemic metabolic function^7,8^, but whether naturally occurring variation in renal nutrient conservation contributes to metabolic disease susceptibility remains unknown. Here, screening eleven genetically divergent mouse strains identified BXD34 mice as strikingly resistant to diet-induced obesity, ectopic lipid accumulation and insulin resistance despite increased caloric intake. Mechanistically, this protection was associated with widespread urinary nutrient loss accompanied by coordinated suppression of renal solute transport, but came at the cost of profound susceptibility to acute kidney injury. Together, these findings identify renal nutrient conservation as an underappreciated determinant of whole-body metabolism and reveal a renometabolic trade-off in which protection from obesity is achieved at the expense of renal resilience.

## Results and Discussion

To investigate natural variation in susceptibility to diet-induced metabolic disease, we exposed eleven genetically divergent mouse strains to chow or high-fat high-sugar (HFHS) diets for six weeks. Ten strains were selected to capture broad phenotypic diversity, with C57BL/6J included as a reference strain.

Metabolic responses to HFHS feeding varied markedly by genetic background. For example, although fat mass increased significantly with HFHS exposure in all strains, glucose tolerance was only impaired in five of eleven, highlighting substantial variation in susceptibility to dietary challenge (**Figure 1A**). BXD34 and CAST were the most resistant strains to obesity, gaining only 3 and 7% adiposity, respectively. Both strains were also resistant to glucose intolerance (**Figure 1B**). Of these two strains, HFHS feeding increased glucose-stimulated insulin levels in CAST but not in BXD34 (**Figure 1C**), distinguishing BXD34 as the most broadly protected strain.

**Figure 1.**
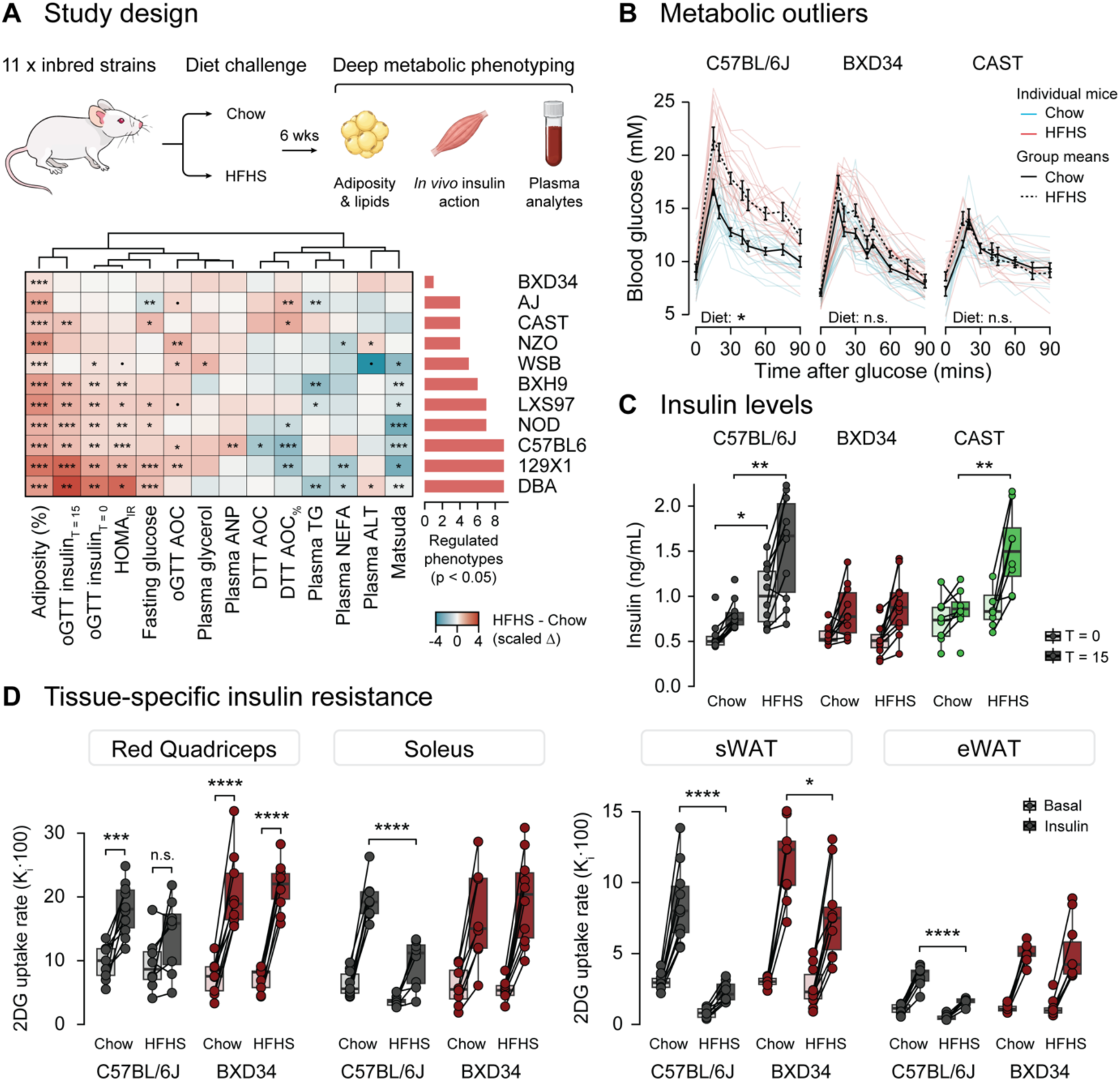
BXD34 mice resist the metabolic effects of high-fat high-sugar feeding. **(A)** Study design for metabolic phenotyping of male mice from eleven genetically divergent inbred mouse strains exposed to chow or HFHS diet for 6 wks (top), and heatmap of diet-induced changes across metabolic phenotypes (bottom). Values represent the delta z-scored phenotypes between chow- and HFHS-fed mice within each strain. Bar plot indicates the number of significantly regulated phenotypes (p < 0.05) within each strain. **(B)** Blood glucose time course during an oral glucose tolerance test (2 g/kg lean mass) in C57BL/6J, BXD34 and CAST mice following chow or HFHS feeding. Data from individual mice are shown as thin coloured lines, and group means as thick black lines. Error bars = SEM. N = 16-22/strain/diet. **(C)** Blood insulin concentrations before and 15 min after glucose administration during the oral glucose tolerance test. **(D)** 2DG uptake into red quadriceps, soleus, subcutaneous white adipose tissue (sWAT) and epididymal white adipose tissue (eWAT) under basal or insulin-stimulated conditions, quantified using the Dual Tracer Test^9^. Minimal data for interpreting systemic insulin action during the DTT is presented in **Extended Data Figure 1**. N = 10/strain/diet. Statistical significance of p < 0.1 is shown as •, p < 0.05 as *, p < 0.01 as **, p < 0.001 as ***, and p < 0.0001 as ****; n.s. = not significant.

To assess insulin action more directly, we used the Dual Tracer Test (DTT) to quantify 2-deoxyglucose (2DG) uptake under basal and insulin-stimulated conditions^9^. HFHS feeding induced insulin resistance in C57BL/6J mice across skeletal muscle (quadriceps and soleus) and adipose tissues (epididymal and subcutaneous white adipose tissue; WAT) (**Figure 1D**). In striking contrast, HFHS-fed BXD34 mice maintained insulin-stimulated 2DG uptake in both skeletal muscles and subcutaneous WAT. Although HFHS feeding impaired insulin-stimulated 2DG uptake in BXD34 epididymal WAT, the magnitude of this impairment was less marked than in C57BL/6J. Because adipose 2DG uptake was normalised to tissue mass, we cannot exclude the possibility that strain differences in HFHS-induced adipocyte hypertrophy influenced these measurements by altering the number of adipocytes represented per unit tissue. Collectively, however, these data demonstrate broad preservation of insulin action across multiple insulin-responsive tissues in BXD34 mice.

To investigate the mechanism underlying the metabolic resilience of BXD34 mice, we exposed a separate cohort to chow or HFHS diet for 5 weeks before metabolic cage analysis. Resistance of BXD34 to diet-induced obesity was strongly replicated (**Figure 2A**).

**Figure 2.**
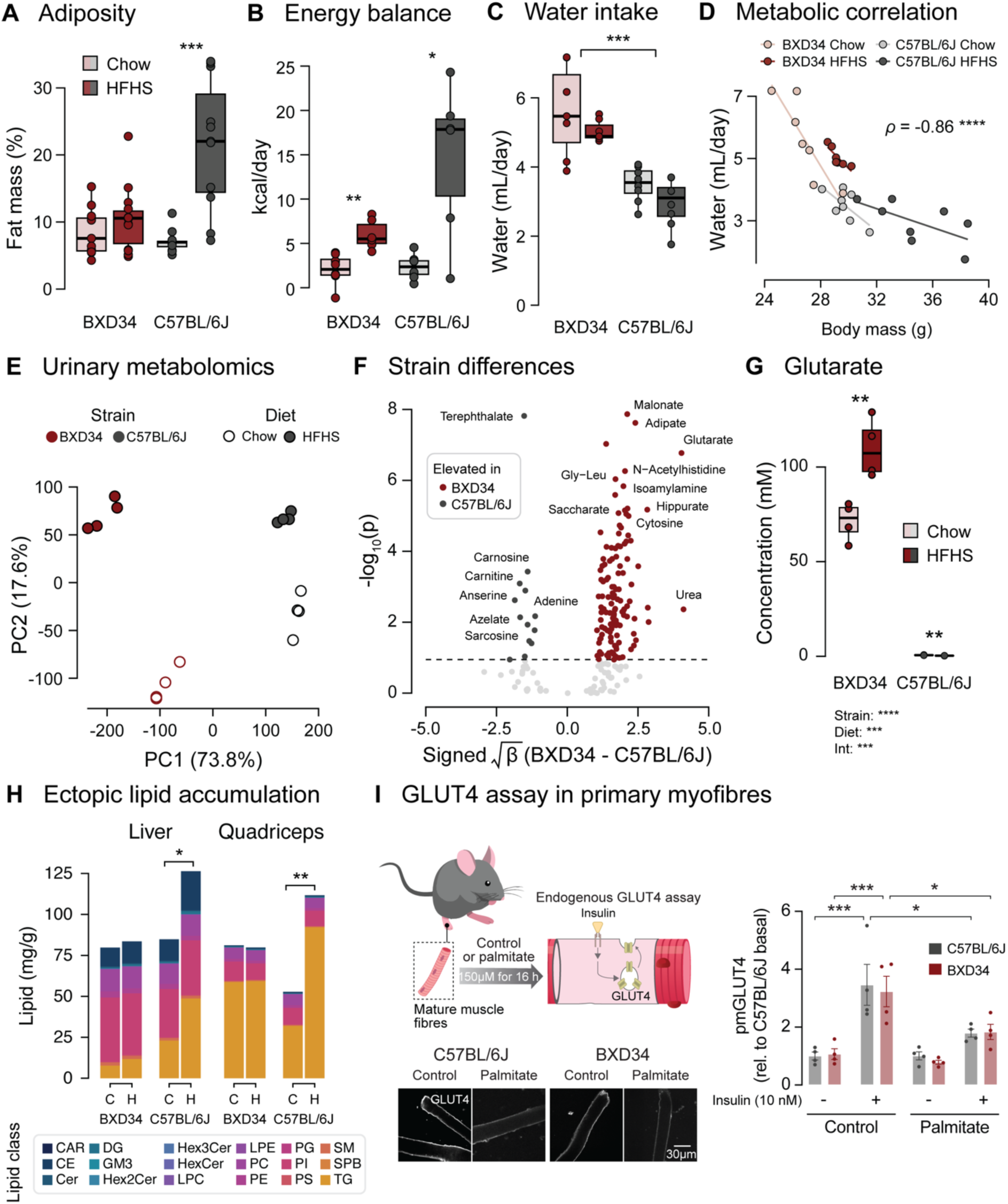
Urinary nutrient loss uncouples caloric excess from metabolic dysfunction. **(A)** Body fat percentages measured immediately before mice went into the metabolic cages. N = 8/strain/diet**. (B)** Energy balance calculated by subtracting energy expenditure from energy intake. **(C)** Total daily water consumption. **(D)** Correlation between body mass and total water intake. **(E)** Principal component analysis of urinary metabolome. N = 4/strain/diet. **(F)** Volcano plot representing the results of a two-way linear model assessing the effects of strain and diet on metabolite abundance. The x-axis represents the signed square root of the main effect of strain. The horizontal dotted line indicates the statistical significance threshold after accounting for multiple testing. **(G)** Urinary glutarate concentrations in chow- and HFHS-fed C57BL/6J and BXD34 mice. **(H)** Total lipid concentrations in liver and quadriceps muscle, coloured by lipid class (255 lipid species across 18 lipid classes). N = 5/strain/diet. Diets: C = chow; H = HFHS. Lipid classes: CAR = acylcarnitine; CE = cholesterol ester; Cer = ceramide; DG = diacylglycerol; GM3 = GM3 ganglioside; Hex2Cer = dihexosylceramide; Hex3Cer = trihexosylceramide; HexCer = hexosylceramide; LPC = lysophosphatidylcholine; LPE = Lysophosphatidylethanolamine; PC = phosphatidylcholine; PE = phosphatidylethanolamine; PG = phosphatidylglycerol; PI = phosphatidylinositol; PS = phosphatidylserine; SM = sphingomyelin; SPB = sphingoid bases; TG = triglycerides. **(I)** Schematic of GLUT4 assay using primary flexor digitorum brevis (FDB) myofibres isolated from adult mice (top left). Representative confocal microscopy images of plasma membrane GLUT4 (pmGLUT4) staining with 10 nM insulin (bottom left). Quantification of pmGLUT4 staining (right). Statistical significance of p < 0.05 is shown as *, p < 0.01 as **, p < 0.001 as ***, and p < 0.0001 as ****.

In C57BL/6J mice, HFHS-induced adiposity was accompanied by increased food intake and positive energy balance (**Figure 2B**; **Extended Data Figure 2A G B**). Strikingly, caloric intake and energy balance were also increased in HFHS-fed BXD34, yet adiposity remained unchanged. This discrepancy was not explained by increased energy expenditure or strain-specific changes in the respiratory exchange ratio (**Extended Data Figure 2B G C**). Instead, BXD34 consumed approximately 1.7-fold more water than C57BL/6J mice (**Figure 2C; Extended Data Figure 2D**), and water intake was strongly inversely correlated with body mass (**Figure 2D**), prompting investigation of whether renal nutrient handling differed between the strains.

Urinary metabolomics revealed profound effects of both strain and diet (**Figure 2E**). More than 70% of metabolomic variation was attributable to strain, compared with approximately 18% to diet. Of 139 metabolites significantly regulated by strain, 126 were elevated in BXD34 (**Figure 2F**). Glutarate showed the largest upregulation, reaching approximately 120- and 350-fold greater abundance in BXD34 compared with C57BL/6J under chow and HFHS conditions, respectively (**Figure 2G**). Together with their increased water consumption, these findings revealed widespread elevation of urinary metabolite loss in BXD34 mice, providing a potential route for the unexplained disposal of dietary nutrients.

We therefore asked whether altered nutrient handling could account for the protection of peripheral tissues from metabolic overload. Lipidomic analysis of liver and quadriceps muscle revealed near-complete protection of BXD34 from HFHS-induced lipid accumulation (**Figure 2H**). To determine whether BXD34 tissues were intrinsically resistant to lipotoxicity, we exposed mature muscle fibres isolated from both strains to 150 µM palmitate *ex vivo* and assessed effects on insulin action. Strikingly, palmitate treatment induced comparable defects in insulin-stimulated GLUT4 trafficking in BXD34 and C57BL/6J muscle fibres (**Figure 2I**). Together, these data indicate that peripheral tissues in BXD34 mice are not intrinsically resistant to lipotoxicity, supporting a model in which their protection *in vivo* instead arises from reduced nutrient exposure associated with elevated urinary nutrient loss.

To assess whether the urinary metabolic profile associated with metabolic health in BXD34 mice was conserved in humans, we analysed data from the OSARTEN cohort^10^, containing 51 urinary metabolites and quantitative metabolic traits across 9,204 individuals. Because many metabolites most strongly upregulated in BXD34, including glutarate, were not measured in this dataset, we derived a human glycaemia-associated signature from metabolites quantified in both datasets. After adjustment for age and biological sex, linear modelling identified nine urinary metabolites significantly associated with fasting blood glucose, a surrogate measure for metabolic health (**Extended Data Figure 2E**). Next, we projected this signature onto the mouse urinary metabolome using single-sample directional gene set analysis (ssdGSA). BXD34 exhibited lower scores, indicating a urinary metabolite profile more closely aligned with lower glycaemia in the human cohort (**Extended Data Figure 2F**). Thus, these findings support a conserved association between urinary metabolite profiles and metabolic health across species.

To identify the molecular programme underlying increased urinary nutrient loss in BXD34, we performed quantitative proteomics across six metabolically relevant tissues from BXD34 and C57BL/6J mice. The kidney exhibited the greatest proteomic divergence between strains, whereas the quadriceps and epididymal white adipose tissue proteomes were largely unaffected by strain (**Figure 3A**). This suggests that protection of these peripheral tissues *in vivo* is driven by factors outside the tissues themselves, such as elevated urinary nutrient loss, rather than by differences in their protein expression.

**Figure 3.**
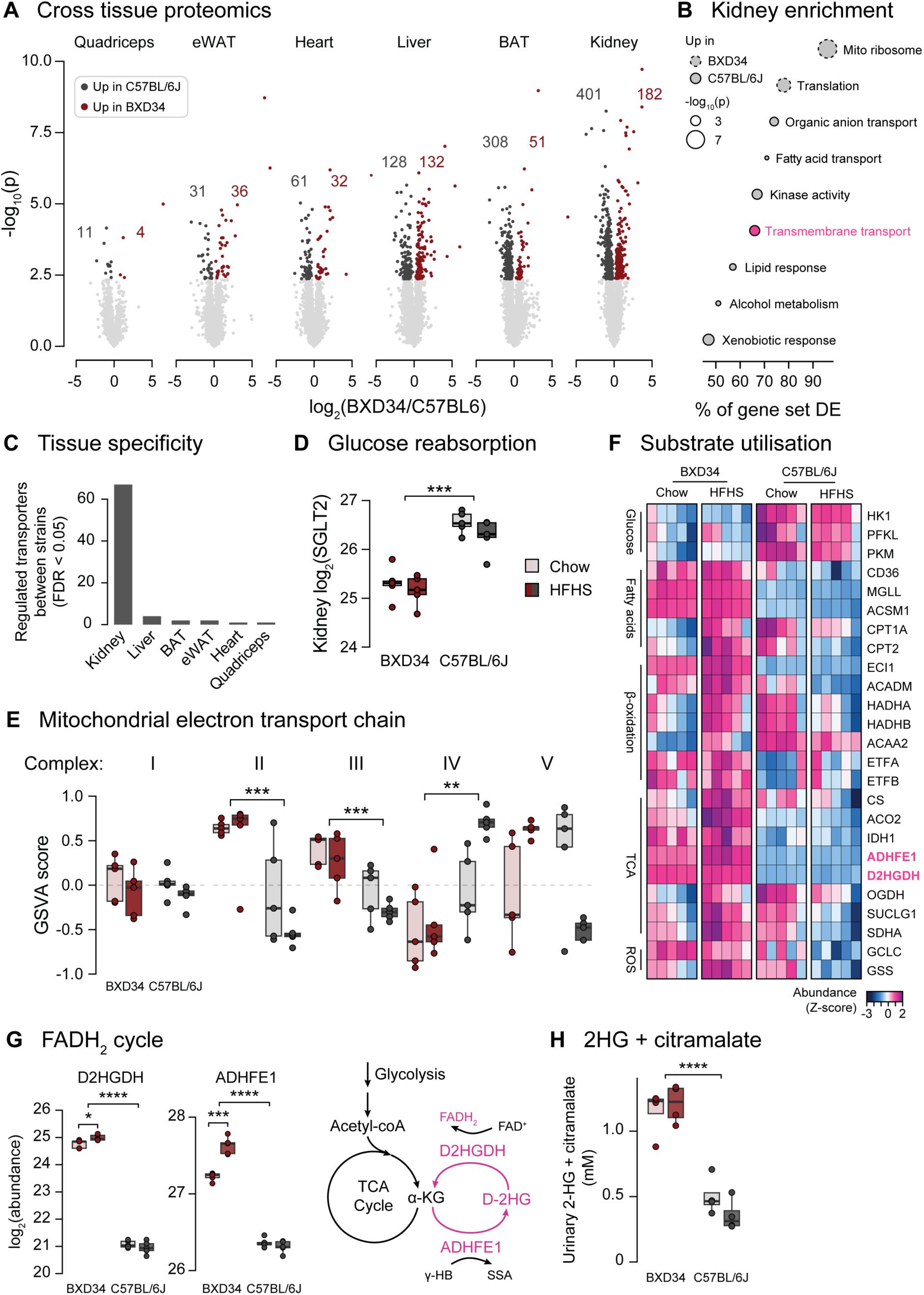
Renal nutrient loss is accompanied by coordinated remodelling of solute transport and fatty acid metabolism. **(A)** Volcano plots showing number of significantly regulated proteins in each tissue under chow conditions (FDR < 0.05; N = 5/strain). BAT = brown adipose tissue. **(B)** Results of gene set enrichment for differentially regulated proteins in the kidney. **(C)** Number of proteins in ‘Transmembrane Transport’ gene ontology that are significantly regulated by strain in each tissue. **(D)** Quantification of SGLT2 abundance in the kidney. **(E)** Gene set variation analysis (GSVA) estimating the relative abundance of mitochondrial electron transport chain complexes in the kidney. Gene sets for each complex were constructed using MitoCarta 3.0^16^. **(F)** Heatmap of z-scored kidney protein abundances for proteins involved in glucose metabolism, fatty acid metabolism, the TCA cycle and antioxidant defence. **(G)** Renal protein abundance of D2HGDH and ADHFE1 (left), and schematic of the proposed FADH_2_-generating cycle (right). **(H)** Combined urinary 2-hydroxyglutarate + citramalate concentration. Statistical significance of p < 0.05 is shown as *, p < 0.01 as **, p < 0.001 as ***, and p < 0.0001 as ****.

Gene set enrichment analysis of the renal proteome identified transmembrane transport as a major altered biological process between strains (**Figure 3B**). Of 67 differentially abundant proteins in this gene ontology, 57 were reduced in BXD34. This regulation was highly specific to the kidney, with fewer than five such proteins varying between strains in any other tissue (**Figure 3C; Extended Data Figure 3A**). Notably, SGLT2 was approximately 2-fold downregulated in BXD34 kidneys (**Figure 3D**). More broadly, approximately 80% of downregulated SLC transporters localised to the plasma membrane (**Extended Data Figure 3B**), and their annotated substrates spanned multiple nutrient classes, including amino acids, carbohydrates, carboxylic acids, lipids and nucleosides (**Extended Data Figure 3C**). This breadth suggests that transporter remodelling in BXD34 mice could affect the reabsorption of a wide range of filtered metabolites. Coordinated suppression of solute transporters beyond SGLT2 therefore provides a candidate molecular mechanism for widespread urinary metabolite loss in BXD34.

To assess the human relevance of this transport programme, we performed phenome-wide enrichment of human genetic associations using *Synteny*^11^, which tests whether a collection of genes is more strongly associated with specific human phenotypes than expected by chance. Genes encoding differentially abundant transmembrane transport proteins were strongly associated with HbA1c (**Extended Data Figure 3D**), with the enrichment signal distributed across multiple genes including *SLC2A2*, *ABCC1* and *SLC16A10*. These findings link the BXD34 renal transport programme to genetically determined variation in human metabolic traits.

Metabolically, BXD34 kidneys exhibited broad remodelling toward increased fatty acid utilisation. This included constitutive changes in the electron transport chain, with increased abundance of complexes II and III (**Figure 3E**), together with increased abundance of proteins involved in fatty acid oxidation and the TCA cycle, and coordinated suppression of glycolytic enzymes including HK1, PFKL and PFKM (**Figure 3F**). Proteins involved in glutathione synthesis were also increased in BXD34 kidneys, consistent with increased oxidative stress accompanying this metabolic shift. BXD34 kidneys displayed further coordinated adaptations predicted to preserve carnitine availability (**Extended Data Figure 3E**), including increased abundance of SLC22A5 (which transports carnitine into cells across the plasma membrane) and SLC25A20 (which transports free and acyl-carnitines into the mitochondria). Despite the widespread increase in urinary metabolite concentrations overall, urinary carnitine was substantially reduced in BXD34 mice (**Extended Data Figure 3F; Figure 2F**), consistent with selective retention of carnitine to support carnitine-dependent fatty acid oxidation.

A particularly striking feature of this metabolic remodelling was the marked upregulation of D2HGDH and ADHFE1 (**Figure 3G**). These enzymes form a metabolic cycle that interconverts α-ketoglutarate and D-2-hydroxyglutarate while coupling redox reactions to FAD/FADH_2_, suggesting engagement of an additional oxidative pathway in BXD34 kidneys. Importantly, 2-hydroxyglutarate (2-HG) exists as two stereoisomers, D-2HG and L-2HG, which are generated and metabolised by distinct enzymatic pathways. In contrast to D-2HG, proteins involved in L-2HG metabolism – including LDHA, MDH2 and L2HGDH – were unchanged between strains, supporting a selective perturbation of D-2HG metabolism. Although our urinary metabolomics approach was unable to resolve total 2HG from citramalate, the combined signal was substantially elevated in BXD34 mice (**Figure 3H**). Given the absence of a canonical pathway for citramalate synthesis in mice, together with increased abundance of D2HGDH and ADHFE1, these findings are consistent with increased D-2HG metabolism in BXD34 kidneys. Considering that D-2HG is elevated in kidney disease^12^ and has established cytotoxic effects^13–15^, these observations raised the possibility that the metabolic adaptations supporting urinary nutrient loss may negatively impact the kidney’s ability to tolerate additional stress.

To determine whether metabolic rewiring in BXD34 kidneys is accompanied by evidence of renal injury, we integrated bulk kidney proteomics with the mouse diabetic kidney disease (DKD) 1M single-cell RNA-sequencing atlas of Wu et al.^17^ (**Figure 4A**). Cell-type composition and gene expression varied markedly between healthy and DKD kidneys in this dataset (**Extended Data Figure 4A G B**), providing cell-type-specific signatures of kidney injury that we used to further interrogate our proteomic data.

**Figure 4.**
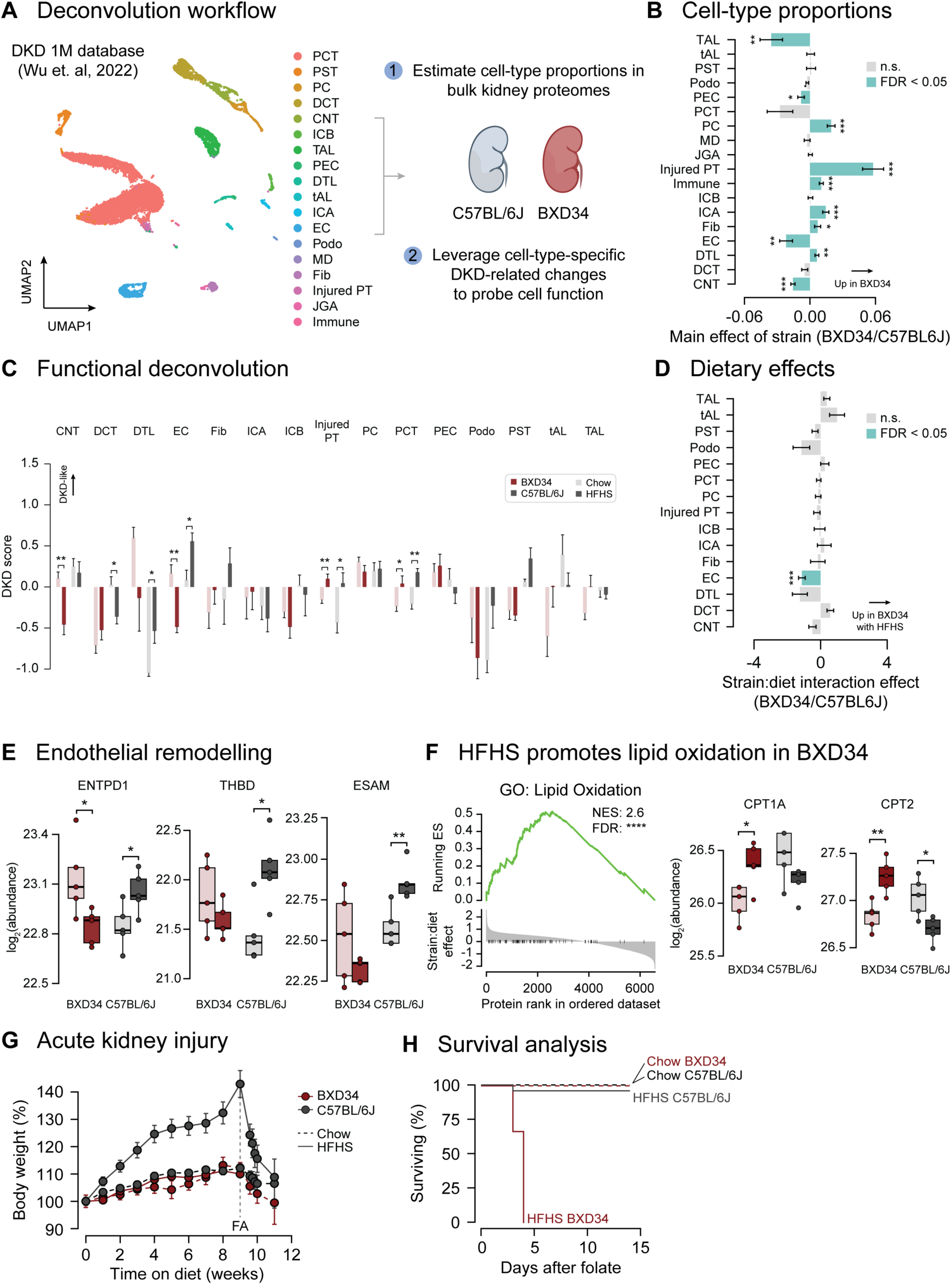
BXD34 mice exhibit latent tubular injury and profound susceptibility to acute kidney injury. **(A)** Workflow for integrating the DKD 1M single-cell RNA-sequencing atlas ^17^ with bulk kidney proteomics. Cell-type identity signatures were used to estimate renal cell-type proportions, while cell-type-specific DKD response signatures were projected onto the kidney proteome to estimate functional remodelling. **(B)** Main effects of strain on renal cell type proportions, determined by two-way linear modelling of the effects of strain and diet. Cell types: TAL = thick ascending limb; tAL = thin ascending limb; PST = proximal straight tubule; Podo = podocyte; PEC = parietal epithelial cell; PCT = proximal convoluted tubule; PC = principal cell; JGA = juxtaglomerular apparatus; ICB = intercalated B; ICA = intercalated A; Fib = fibroblast; EC = endothelial cell; DTL = descending thin limb; DCT = distal convoluted tubule; CNT = connecting tubule. **(C)** Cell-type-specific DKD scores across chow- and HFHS-fed C57BL/6J and BXD34 mice. Higher values indicate greater similarity to the DKD-associated state for the corresponding cell type. **(D)** Strain-by-diet interaction effects for cell-type-specific DKD scores. Positive values indicate a greater HFHS-induced increase in BXD34 relative to C57BL/6J. **(E)** Renal abundance of endothelial proteins ENTPD1, THBD and ESAM. **(F)** Gene set enrichment analysis of strain-by-diet interaction effects demonstrating enrichment of lipid oxidation pathways, with representative renal abundance of CPT1A and CPT2 (right). **(G)** Mean body weight during dietary intervention and following folic acid administration. FA indicates the timing of folic acid administration. N = 16/strain/diet. Error bars represent SEM. **(H)** Survival following folic acid-induced acute kidney injury in chow- and HFHS-fed C57BL/6J and BXD34 mice. Statistical significance of p < 0.05 is shown as *, p < 0.01 as **, and p < 0.001 as ***.

Cell-type deconvolution revealed marked differences in renal cellular composition between strains. The BXD34 genotype most strongly associated with an increased inferred proportion of injured proximal tubule (PT) cells (**Figure 4B**), consistent with a pre-existing injury-associated tubular state. HFHS feeding produced relatively modest changes in inferred cell-type composition (**Extended Data Figure 4C**). In contrast, analysis using cell-type-specific DKD signatures revealed substantial functional remodelling (**Figure 4C**). Injured PT cells adopted a more DKD-like signature in both strains, while endothelial cells (EC) showed divergent responses to HFHS feeding between BXD34 and C57BL/6J (**Figure 4C G D**).

The endothelial response was particularly striking. ENTPD1, an endothelial ectonucleotidase implicated in vascular protection and recovery after renal injury^18,19^, decreased with HFHS feeding in BXD34 but increased in C57BL/6J. C57BL/6J also showed increased THBD and ESAM – proteins associated with endothelial protection and vascular integrity^20–23^ – whereas these responses were absent in BXD34 (**Figure 4E**). These findings are consistent with an adaptive endothelial response to HFHS diet in C57BL/6J that is absent or blunted in BXD34, potentially limiting the capacity of BXD34 kidneys to tolerate additional injury. In parallel, proteins exhibiting significant strain-by-diet interactions were strongly enriched for fatty acid oxidation pathways (**Figure 4F**), consistent with greater metabolic remodelling of the BXD34 kidney during HFHS feeding. Together, these findings suggest that the metabolic adaptations observed in BXD34 mice might render the kidney less resilient to additional stress.

We therefore challenged chow- and HFHS-fed C57BL/6J and BXD34 with folic acid-induced acute kidney injury, a well-established model of tubular nephrotoxicity^24^. To avoid confounding injury severity due to differences in body mass, mice received a fixed dose of 7.5 mg of folic acid, equivalent to 250 mg/kg in a 30 g mouse.

All groups exhibited significant weight loss and renal injury following folic acid administration (**Figure 4G; Extended Data Figure 4D**). Strikingly, folic acid administration was lethal in 100% of HFHS-fed BXD34 mice, compared with 12% of HFHS-fed C57BL/6J mice and no lethality in either chow-fed group (**Figure 4H**). Although the increase in lethality in HFHS-fed C57BL/6J was not statistically significant, histological assessment and blood urea nitrogen measurements confirmed greater folic acid-induced renal injury in C57BL/6J (**Extended Data Figure 4E G F**), consistent with previous reports^25,26^.

Together, these data reveal a striking renometabolic trade-off: the renal adaptations associated with metabolic protection against diet-induced metabolic disease are accompanied by profound vulnerability to kidney injury under metabolic stress.

SGLT2 inhibitors have been used to treat type 2 diabetes for more than a decade^27^, demonstrating that altering renal nutrient reabsorption can produce substantial metabolic and cardiovascular benefit. Yet, whether naturally occurring variation in renal nutrient conservation contributes to susceptibility to metabolic disease remains poorly understood. Here, we identify renal nutrient loss as an underappreciated determinant of susceptibility to diet-induced obesity and metabolic dysfunction. More broadly, our findings illustrate how genetic diversity can uncover physiological states that redistribute disease susceptibility between organs, rather than simply conferring uniform disease resistance^28^.

Screening genetically divergent mouse strains identified BXD34 as strikingly resistant to HFHS-induced obesity and insulin resistance despite increased caloric intake. This paradox was associated with widespread urinary metabolite loss and coordinated suppression of renal solute transport, while peripheral tissues remained intrinsically susceptible to lipotoxicity when challenged *ex vivo*. Consistent with the broader relevance of renal nutrient handling, genetic variation in renal solute transport machinery was strongly associated with HbA1c in humans, while urinary metabolite profiles were independently associated with glycaemia. These findings extend the conventional framework of energy balance by identifying renal nutrient excretion as a critical determinant of how much dietary energy is ultimately retained by the body.

BXD34 mice represent an extreme example of renal nutrient loss. Whereas SGLT2 inhibitors selectively promote urinary glucose excretion and typically produce only modest weight loss^29^, BXD34 mice exhibited broad suppression of renal solute transport and near-complete protection from HFHS-induced adiposity. Familial renal glucosuria (FRG), caused by loss-of-function variants in *SLC5A2*, provides a naturally occurring human analogue of chronically reduced renal glucose conservation, with urinary glucose excretion reaching up to 160 g/day in severe cases^30,31^. Altered glucose handling in FRG can also perturb the reabsorption of other proximal tubular substrates^32,33^ and has been associated with reduced obesity risk^34^. Importantly, however, FRG is not typically associated with tubular injury^31,35,36^. Substantial renal nutrient loss, therefore, does not inevitably compromise renal function. This suggests that the renal vulnerability of BXD34 arises not simply from nutrient wasting itself, but from the broader physiological adaptations associated with its transport phenotype.

Our data point to several candidate components of this adaptation. Renal solute transport is energetically demanding, and changes in proximal tubular transport can profoundly alter oxygen consumption and substrate metabolism^37,38^. Reduced solute transport in BXD34 mice was accompanied by extensive metabolic remodelling, including fatty acid oxidation and antioxidant machinery, altered composition of the electron transport chain, and selective carnitine retention. D2HGDH and ADHFE1 were also markedly increased, implicating altered D-2HG metabolism and FAD-dependent redox reactions. Given that D-2HG can exert cytotoxic effects and has been linked to oxidative stress and kidney disease^12–15^, this pathway represents one potential contributor to renal vulnerability in BXD34 mice. However, neither increased D-2HG flux nor a causal role for D-2HG is established by our data. Rather, the broader proteomic signature indicates that BXD34 kidneys occupy a substantially remodelled metabolic and redox state that may reduce their capacity to accommodate additional metabolic stress, as observed upon HFHS feeding.

Importantly, evidence of renal vulnerability extended beyond the above metabolic changes. BXD34 kidneys exhibited an increased inferred abundance of injured proximal tubule cells even under chow-fed conditions, while HFHS feeding induced additional DKD-like functional remodelling. Most notably, endothelial cell responses to HFHS feeding differed markedly between strains: C57BL/6J kidneys increased ENTPD1, THBD, and ESAM – associated with endothelial protection and vascular integrity^20–23^ – whereas these responses were absent or reversed in BXD34 mice. These observations are consistent with an adaptive endothelial response in C57BL/6J that is diminished in BXD34. We therefore propose that renal vulnerability in BXD34 reflects the combined effects of pre-existing tubular injury, altered metabolic and redox homeostasis, and impaired endothelial adaptation, rather than a single lesion. This latent vulnerability was exposed by folic acid-induced acute kidney injury, which was lethal in all HFHS-fed BXD34 mice but caused substantially less mortality in all other groups.

Together, these findings expose a renometabolic trade-off in which renal adaptations that limit systemic nutrient retention protect against obesity and insulin resistance at the expense of reduced renal resilience. The therapeutic question raised by this trade-off, therefore, is not simply whether renal nutrient loss can be increased, but whether its systemic metabolic benefits can be uncoupled from the adaptations that compromise the kidney’s ability to withstand additional stress. The tolerance of substantial nutrient loss in FRG suggests that such uncoupling may be possible. Defining which components of the BXD34 programme confer systemic metabolic protection and which impose the renal cost could reveal novel strategies to reproduce the systemic benefits of urinary nutrient loss without sacrificing kidney health.

## Methods

### Mouse handling

129X1/SvJ (129X1), BXH9/TyJ (BXH9), BXD34/TyJ (BXD34), CAST/EiJ (CAST), ILSXISS97/TejJ (LXS97), NZO/HILtJ (NZO) and WSB/EiJ (WSB) inbred mouse strains were obtained from Australian BioResources (NSW, Australia). C57BL/6J, DBA/2J (DBA), A/J and NOD/ShiLtJ (NOD) inbred mouse strains were obtained from Animal Resources Centre (WA, Australia). Experiments were performed in accordance with NHMRC (Australia) guidelines under the approval of The University of Sydney Animal Ethics Committee. Mice were monitored twice/wk and weighed weekly. Fat and lean mass measures were acquired via EchoMRI-900 (EchoMRI Corporation).

All experiments were performed in male mice. Mice were maintained at 23°C on a 12 h light/dark cycle and given *ad libitum* access to food and water in individually ventilated cages with a density of 2-5 mice/cage. Mice were acclimatised for 1 wk prior to experimentation. Mice were enrolled in studies between 12-14 wks of age and were randomised to either a standard laboratory chow diet containing 13% calories from fat, 65% calories from carbohydrate and 22% calories from protein (‘Irradiated Rat and Mouse Diet’, Specialty Feeds), or a high-fat high-sugar (HFHS) diet made in-house containing 45% calories from fat, 35% calories from carbohydrate and 20% calories from protein; manufactured to closely resemble D12451 (Research Diets). Specifically, the HFHS diet contained 23% w/w casein, 0.3% w/w methionine, 2% w/w gelatine, 20.2% w/w sucrose, 17% w/w corn starch, 5% w/w bran, 3% w/w safflower oil, 22% w/w lard, 5.8% w/w AIN-93 mineral mix (MP Biomedicals), 0.4% w/w choline bitartrate and 1.3% w/w AIN-93 vitamin mix (MP Biomedicals).

Urine was collected in 5 mL Eppendorf tubes by scruffing mice and applying gentle pressure to the lower abdomen. Plasma was collected by making a small incision in the tail and collecting whole blood into an EDTA-coated tube on ice. Whole blood samples were spun at 2,000 g for 10 min at 4 °C and the plasma supernatant transferred to a clean tube. All biofluids were stored at - 80 °C until analysis. Tissues were collected from mice following cervical dislocation. Tissues were rapidly excised, snap-frozen in liquid nitrogen and stored at -80 °C until further processing.

Samples were randomised and identified only by numerical IDs during experimental processing and analysis.

### Oral glucose tolerance test

Mice fasted for 6 h (0700 – 1300 h) were dosed via oral gavage of 25% glucose in water at 2 g/kg lean mass. Blood glucose concentration was measured directly from tail whole blood via a glucose monitor (Roche Diabetes Care) at 0, 20, 30, 40, 60 and 75 and 90 min post glucose administration. The baseline-corrected area of the curve (AOC) was calculated as:

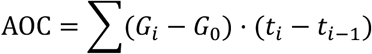

Where G and t equal the glucose concentration and time at the *i^t^*^ℎ^ blood sample, respectively; and *G*_0_ is the fasting glucose concentration. Differences in AOC between mice reflect altered glucose tolerance independent of differences in fasting glucose concentrations.

Blood insulin concentrations were measured by collecting 5 µL of blood from tail veins at 0 and 15 min directly into an Insulin Mouse Ultra-Sensitive ELISA kit (Crystal Chem USA, catalogue #90080) containing sample diluent on ice. The assay was then performed according to the manufacturer’s protocol, using an insulin standard curve to determine sample insulin concentrations.

### Dual Tracer Test

Dual Tracer Tests were performed as described previously^9^. Briefly, mice were fasted for 2 h, anaesthetised with sodium pentobarbital (65 mg/kg) and kept euthermic by using foil and heat pads. A first retro-orbital injection of 2.5 µCi ^14^C-2DG in plasma replacement (B. Braun, catalogue #FV41513) was administered, with blood collected at 2, 15 and 30 min. At 40 min, mice received 5 μCi ^3^H-2DG plus insulin (0.75 U/kg lean mass), followed by blood sampling at 2, 15 and 30 min. Mice were euthanised by cervical dislocation at 80 min and tissues rapidly excised, snap-frozen and stored at -80 °C.

Whole body tracer clearance was determined from the exponential decay of the circulating 2DG tracers. Tissues were powdered, lysed in 1% Triton X-100 (Sigma-Aldrich, catalogue #X100), and glucose uptake normalised to tissue mass. Non-phosphorylated 2DG was separated using AG 1-X8 anion exchange resin chloride form (Bio-Rad, catalogue #1401441), and tracer levels measured by liquid scintillation counting. Phosphorylated 2DG was calculated as the total minus non-phosphorylated counts. For tissues other than the heart, ^14^C-2DG-6P accumulation was used as an estimate of basal glucose uptake, as validated previously^9^.

Tissue-specific uptake rates were calculated using the method of Hom et al.^39^, adjusting for initial dose, tracer kinetics, and exposure time (80 min basal, 40 min insulin-stimulated). Insulin-stimulated uptake was determined by subtracting basal from insulin-stimulated 2DG uptake.

### Indirect calorimetry

Metabolic cage experiments were performed using a PhenoMaster NG metabolic cage system (TSE Systems) after 5 wks exposure to chow or a HFHS diet. Mice were acclimatised to the PhenoMaster system for 5 days prior to the 48 h recording period. Oxygen intake (VO2) and carbon dioxide output (VCO2) were measured every 20 min to determine the respiratory exchange ratio (RER) and calculate energy expenditure (EE). Infrared beams were used to measure activity by recording beam breaks. Sleep cycles were deduced from extended periods of no beam breaks, suggesting inactivity. Food and water consumption were also monitored continuously. Mice were returned to their home cages with cage-mates at the end of the recording period. Energy balance was calculated by subtracting energy expenditure from total energy intake.

Data analysis was performed in line with current best practices^40^. For summarised quantification, parameters were aggregated across a 48 h recording period as well as within light and dark phases. Real-time recording traces are visualised as running hourly averages. All statistical analyses were performed using raw data.

### Urinary metabolomics

For metabolite extraction, 10 µL urine was mixed with 90 µL MS-grade H_2_O containing 200 µM of each internal standard (D-camphor-10-sulfonic acid (CSA), 2-(N-morpholino)-ethanesulfonic acid (MES), methionine sulfone, trimesic acid, and 3-aminopyrrolidine). 90 μL of the solutions were transferred to 5 kDa centrifugal ultrafiltration tubes (Human Metabolome Technologies) and centrifuged for 1 h at 9,100 g at 20 °C. Cationic metabolomic analysis was performed using undiluted filtrates, whereas anionic metabolic analysis was performed using a 1:1 dilution of filtrate to MilliQ water.

Capillary electrophoresis time-of-flight mass spectrometry (CE-TOFMS) experiments were performed using an Agilent 7100 CE capillary electrophoresis system, Agilent 6230 LC/MSD TOF mass spectrometry system, Agilent 1100 series binary HPLC pump, Agilent CE-MS adaptor (catalogue #G1603A) and Agilent CE-ESI-MS sprayer kit (catalogue #G1607A). For anionic metabolite analysis, the original Agilent stainless ESI needle was replaced with an Agilent platinum ESI needle (catalogue #G7100-60041)^41^. System control and data acquisition were performed using an Agilent MassHunter Workstation, and data analysis was performed using the proprietary in-house Keio MasterHands software^42–46^.

For cationic metabolite analysis, separations were carried out in a fused silica capillary (50 µm x 97 cm) filled with 1 M formic acid as the electrolyte^41,46^. Approximately 5 nL of sample solution were injected at 50 mbar for 5 s, and then electrophoresis was performed with 30 kV of voltage. The capillary temperature was maintained at 20 °C and the sample tray was cooled below 5 °C. Methanol-water (50% v/v) containing 0.01 µM Hexakis-(2,2-difluoroethoxy)-phosphazene was delivered as the sheath liquid at 10 µL/min. Electrospray ionization (ESI)-time-of-flight mass spectrometry (TOFMS) was conducted in the positive ion mode and the capillary voltage was set at 4,000 V. A flow rate of heated dry nitrogen gas (heater temperature 300 °C) was maintained at 7 psig. In TOFMS, the fragmentor-, skimmer-, and Oct RFV voltage was set at 75 V, 50 V, and 500 V, respectively. Automatic recalibration of each acquired spectrum was performed using reference masses of reference standards. The ^13^C isotopic ion of a protonated methanol dimer ([2MeOH+H]+, m/z 66.0631) and Hexakis-(2,2-difluoroethoxy)-phosphazene ([M+H]+, m/z 622.0290) provided the lock mass for exact mass measurements^47,48^.

For anionic metabolite analysis, a commercially available COSMO(+) capillary (50 µm x 102 cm) (Nacalai Tesque, catalogue #07584-44) was used with a 50 mM ammonium acetate solution (pH 8.5) as the electrolyte^41,46^. Sample solution (30 nL) was injected at 50 mbar for 30 sec, and then electrophoresis was performed with -30 kV. Ammonium acetate (5 mM) in 50% methanol-water (v/v) containing 0.01 μM Hexakis-(2,2-difluoroethoxy)-phosphazene was delivered as the sheath liquid at 10 µL/min. ESI-TOFMS was conducted in the negative ion mode and the capillary voltage was set to 3,500 V. For TOFMS, the fragmentor-, skimmer-, and Oct RFV voltage was set at 100 V, 50 V, and 500 V, respectively. Automatic recalibration of each acquired spectrum was performed using reference masses of reference standards. The ^13^C isotopic ion of a deprotonated acetic acid dimer ([2CH3COOH-H]-, m/z 120.0384), and Hexakis + deprotonated acetic acid (m/z 680.03554) provided the lock mass for exact mass measurements^47,48^.

### Tissue proteomics

Frozen tissues were homogenised using a liquid nitrogen cooled mortar and pestle before being weighed into a clean tube on dry ice. Approximately 20 mg of powdered tissues were lysed in 600 µL 4% sodium deoxycholate (SDC; Thermo Fisher, catalogue #B20759) in 100 mM Tris pH 8.5 buffer by boiling at 95 °C for 10 min with continuous vortexing in a thermomixer. Lysates were sonicated for 30 s using a tip-probe sonicator and centrifuged for 10 min at 20,000 g to remove insoluble material. Supernatants were transferred to a clean tube and cleaned by high-volume chloroform-methanol precipitation. 1600 µL MS-grade methanol, 800 µL MS-grade chloroform and 800 µL MS-grade H_2_O were added in sequence with brief vortexing between each addition. Samples were centrifuged for 5 min at 2,000 g to achieve phase separation. The upper aqueous phase was carefully removed, and the protein pellet washed with 2400 µL MS-grade methanol. Samples were centrifuged for 5 min at 2,000 g to pellet the protein and the entire supernatant was removed. The protein pellet was resuspended in 200 µL 4% sodium deoxycholate in 100 mM Tris pH 8.5.

Kidney and quadriceps samples underwent an additional acetone precipitation to remove residual contaminants. 1800 µL -30 °C MS-grade acetone was added to resuspended protein lysates and incubated at -30 °C overnight. Samples were sonicated for 30 s with a tip probe sonicator and centrifuged for 15 min at 2,500 g at 4 °C. The supernatant was removed and replaced with 800 µL -30 °C 80% acetone. Samples were again sonicated for 30 s and returned to -30 °C for 2 h. After this, samples were centrifuged for 15 min at 2,500 g at 4 °C and supernatant discarded. Tubes were left to air dry for 10 min before resuspension of the protein pellet in 200 µL 4% sodium deoxycholate in 100 mM Tris pH 8.5.

Protein concentration was determined by BCA assay. 10 µg of protein was aliquoted into 1.5 mL centrifuge tubes and adjusted to 1% SDC with MS-grade water before reduction/alkylation buffer (10 mM TCEP, 40 mM 2-chloroacetamide) was added and the samples heated for 15 min at 60 °C. Once cooled to RT, 0.2 µg trypsin and 0.2 µg LysC were added to each sample and incubated overnight (18 h) at 37 °C. An equal volume of 1% trifluoroacetic acid (TFA) in ethyl acetate was added to each sample to stop digestion and remove SDC.

Samples were prepared for LC-MS/MS analysis by StageTip clean up using SDB-RPS solid phase extraction material^49^. Briefly, 3 layers of SDB-RPS material were packed into 200 µL tips and samples were loaded onto StageTips by centrifugation at 1,000 g for 5 min. StageTips were washed with subsequent spins at 1,000 g for 5 min with 100 µL 1% TFA in ethyl acetate, then 1% TFA in isopropanol, and 0.2% TFA in 5% acetonitrile (ACN). Samples were eluted by addition of 100 µL 60% ACN with 5% NH_4_OH. Samples were dried by vacuum centrifugation and reconstituted in 40 µL 5% formic acid.

150 ng of peptides were injected onto an in-house assembled LC column (75 µm x 45 cm) packed with 1.9 µm C18 ReproSil Pur AQ particles (Dr. Maisch HPLC GmbH). Peptides were separated by gradient elution using a Vanquish Neo UHPLC System (Thermo Fisher) with a combination of buffer A (0.1% formic acid in water) and buffer B (0.1 % formic acid in 80% ACN). Samples were loaded at a constant pressure of 750 bar with 97% buffer A, followed by a gradual increment to 40% B over 66 min at 0.3 µL/min, and then to 98% B over 4 min. The column was washed for 5 min with 98% B at 0.5 μL/min before returning to 97% A in preparation for the next sample. Column temperature was maintained at 60 °C using a Sonation column oven.

Eluting peptides were coupled to tandem mass spectrometry (MS/MS) using a Q-Exactive HF-X mass spectrometer (Thermo Fisher Scientific). Peptides were ionised by electrospray under a voltage of 2.4 kV, with a transfer capillary temperature of 300 °C. MS1 spectra were collected between mass/charge (m/z) 350 and 1650 (resolution = 120,000). Fragmentation spectra were collected using data-independent acquisition (DIA), with variable-width isolation windows ranging from 27 to 589 m/z units in size (**Table 1**). Peptides in each isolation window were subjected to higher-energy collisional dissociation (collision energy = 25%) to obtain fragment ions. MS2 spectra were collected between m/z 300 and 2000 (resolution = 30,000, AGC target = 3e^6^, maximum injection time = automatic).

**Table 1.** Variable width isolation windows for mass spectrometry method.

| DIA Window | Min | Max | m/z centre | Window width (m/z) |
| --- | --- | --- | --- | --- |
| 1 | 350 | 394 | 372.0 | 44 |
| 2 | 393 | 424 | 408.5 | 31 |
| 3 | 423 | 452 | 437.5 | 29 |
| 4 | 451 | 478 | 464.5 | 27 |
| 5 | 477 | 504 | 490.5 | 27 |
| 6 | 503 | 529 | 516.0 | 26 |
| 7 | 528 | 555 | 541.5 | 27 |
| 8 | 554 | 581 | 567.5 | 27 |
| 9 | 580 | 608 | 594.0 | 28 |
| 10 | 607 | 635 | 621.0 | 28 |
| 11 | 634 | 663 | 648.5 | 29 |
| 12 | 662 | 693 | 677.5 | 31 |
| 13 | 692 | 725 | 708.5 | 33 |
| 14 | 724 | 759 | 741.5 | 35 |
| 15 | 758 | 798 | 778.0 | 40 |
| 16 | 797 | 841 | 819.0 | 44 |
| 17 | 840 | 892 | 866.0 | 52 |
| 18 | 891 | 959 | 925.0 | 68 |
| 19 | 958 | 1062 | 1010.0 | 104 |
| 20 | 1061 | 1650 | 1355.5 | 589 |

Proteomics raw data files were searched using DIA-NN version 1.9.2 using library-free FASTA searches against the reviewed UniProt mouse proteome with deep learning enabled^50^. The protease was set to Trypsin/P with 1 missed cleavage, and N-term Methionine excision, carbamidomethylation and Methionine oxidation options were also set to on. Peptide length was set to 7-30, precursor range 350-1650 m/z, fragment range 300-2000 m/z and FDR set to 1%. To reduce false-positive protein identifications, match between runs was only performed for samples from the same tissue. Protein intensities were log_2_-transformed and median-normalised for all analyses.

### Tissue lipidomics

Approximately 15 mg powdered tissue was lysed in 225 µL MS-grade methanol (Fisher Chemical, catalogue #A456) by vortexing for 20 s, incubating on dry ice for 1 min and then sonicating with a tip-probe sonicator for 30 s. The freeze-thaw-vortex cycle was repeated another 3 times to ensure complete tissue rupture. Next, 750 µL MS-grade methyl-tert-butyl ether (MTBE; Thermo Fisher, catalogue #M-4496-17) was added to each sample and incubated for 1 h at RT while vortexing at 750 rpm using a ThermoMixer C (Eppendorf, catalogue #5382000066). Separation between aqueous and lipid phases was induced by the addition of 190 µL MS-grade H_2_O (Fisher Chemical, catalogue #W64) followed by brief vortexing and centrifugation for 5 min at 10,000 g. MTBE lipid extracts (600 µL) were collected from the upper phase into a clean tube and dried using a speed-vacuum concentrator and reconstituted in 500 µL chilled butanol:methanol (1:1 v/v) for LC-MS/MS analysis. A sample made of pooled aliquots of lipid extracts was used as a quality control sample and injected every 10 study samples.

Analysis was performed using an Agilent UHPLC 1290 Infinity II liquid chromatography system connected to an Agilent QqQ 6495C.

An Agilent Zorbax RRHD Eclipse Plus C18 column (2.1 × 50 mm, 1.8 µm) was used for the RPLC separation. The mobile phases A (60% H_2_O and 40% ACN with 10 mM ammonium formate) and B (10% ACN and 90% isopropanol with 10 mM ammonium formate) were used for the chromatographic separation. The following gradient was applied: 0-2 min, 20-60% B; 2-12 min, 60-100% B; 12-14 min, 100% B; 14.01-15.8 min, 20% B. The oven temperature was maintained at 40 °C. Flow rate was set at 0.4 mL/min and the sample injection volume was 1 µL.

The positive ionisation spray voltage and nozzle voltage were set at 3,000 V and 1,000 V, respectively. The drying gas and sheath gas temperatures were both maintained at 250 °C. The drying gas and sheath gas flow rates were 14 L/min and 11 L/min, respectively. The nebulizer nitrogen gas flow rate was set at 35 psi. The iFunnel high- and low-pressure RF were 150 V and 60 V, respectively. Targeted analysis was performed in Dynamic MRM positive ion mode.

The acquired MS data were analysed using the Agilent MassHunter software version B.10.00. The signal-to-noise ratios (S/N) were calculated using the raw peak areas in study samples and blanks (BLK). Lipids that had CV > 20% in the QC samples, QC/BLK < 10 and did not show a linear behaviour (R^2^ < 0.8) in dilution curves were excluded from the analysis. Internal standards were used to normalize the raw peak areas in the corresponding lipid class and concentrations were further normalized to the tissue weight in the original sample.

### Flexor digitorum brevis GLUT4 assay

Muscle fibres were isolated as described previously^51^. Briefly, flexor digitorum brevis (FDB) muscles were carefully dissected and incubated in a solution containing 4 mg/mL collagenase type I (Worthington, catalogue #9001-12-1) in minimum essential medium (Gibco, catalogue #A10489010) supplemented with 10% FBS (Sigma Aldrich, catalogue #F8192) for 4 h at 37 °C in a 5% CO_2_ incubator. Following enzymatic digestion, fibres were mechanically dissociated using fire-polished glass pipettes with progressively smaller diameters. Isolated fibres were plated into 96-well plates coated with laminin (Thermo Fisher, catalogue #23017015) and maintained in minimal essential medium supplemented with 10% FBS, 50 µg/mL gentamicin (Gibco, catalogue#10131035), 100 µg/mL streptomycin, and 100 IU/mL penicillin (Gibco, catalogue #15140122). Fibres were incubated at 37 °C with 5% CO_2_.

To induce *in vitro* insulin resistance, cells were treated for 16 h with 150 µM palmitate-BSA or ethanol-BSA as a control^52^. Palmitate was complexed with BSA by dissolving the fatty acid in 100% ethanol and then diluting 25-fold in a 10.5% fatty acid-free BSA solution. These stock solutions were further diluted in culture media to achieve a final concentration of 150 µM. Palmitate treatments were initiated 3 h after cells were seeded into experimental plates.

Endogenous GLUT4 assays were performed as described previously^52^. Fibres were washed with warm PBS and serum-starved for 2 h before being stimulated with 10 nM insulin for 20 min. Fibres were then washed three times with ice-cold PBS while on ice. Extracellular GLUT4 staining was achieved by incubating fibres in a solution containing 10% horse serum and 10 µg/mL exofacial GLUT4 antibodies LM048 and LM052, provided by Joseph Rucker (Integral Molecular)^53^ for 20 min on ice. Fibres were then fixed with 4% paraformaldehyde (Electron Microscopy Sciences, catalogue #15710) for 5 min on ice and then 20 min at room temperature. Following fixation, cells were washed twice with PBS and incubated with 50 mM glycine in PBS for 5 min at RT. Next, a secondary antibody (AF647-conjugated goat anti-human; Jackson ImmunoResearch, catalogue #109-607-003) diluted 1:200 in 10% horse serum was incubated for 1 h at room temperature. The secondary antibody was removed, and cells were washed three times with degassed PBS, prior to the addition of 100 µL/well degassed imaging buffer (2.5% 1,4-diazabicyclo[2.2.2]octane and 10% glycerol, pH 7.8). Solutions containing primary or secondary antibodies were filtered through a 0.22-µm syringe filter (Millipore, catalogue #SLGP033NB) prior to use.

Images were taken in a single Z-plane at the midsection of most muscle fibres using a conventional confocal system with 2 x 2 pixel binning and an 8 x 8 montage to capture the entire well. Fluorescence at the plasma membrane was quantified semi-automatically. Healthy fibres were manually selected using Fiji, and CellProfiler^54^ was used to measure PM fluorescence. Muscle fibres were selected using bright-field microscopy based solely on morphology, thereby avoiding any bias related to insulin response. Fibres were chosen if they were straight and displayed clear striated structures. A global threshold based on 488-nm autofluorescence created a whole-fibre mask, followed by a 4-pixel-wide perimeter mask for quantifying GLUT4 fluorescence intensity at 647 nm.

### Acute kidney injury

To induce acute kidney injury, mice were administered a single dose of 7.5 mg folic acid (Sigma Aldrich, catalogue #F7876) after 8 wks exposure to a chow or HFHS diet by intraperitoneal injection. This dosing scheme is equivalent to the established dose of 250 mg/kg^24^ for a 30 g mouse, and was selected to maintain comparable injury stimulus between chow and HFHS-fed mice of varying body weights.

Urine, plasma and kidney samples were taken from independent terminal cohorts at T = 0, 4, 7 and 14 days post folic acid administration. The right kidney was excised, bisected longitudinally and fixed in 4% paraformaldehyde for 48 h for histological analysis. Fixed kidneys were sectioned (3 µm) and then stained using the periodic acid-Schiff method to identify glomerular and tubular basement membranes. Slides were scanned to digital images using an Aperio ScanScope at 20x magnification. The left kidney was dissected to carefully remove the renal capsule and any adipose tissue encompassing the kidney, before snap freezing in liquid nitrogen and storing at - 80 °C for subsequent analysis.

Blood urea nitrogen assays were performed using a colourimetric kit (Thermo Fisher, catalogue #EIABUN). Plasma was diluted 1:20 with sample diluent, and the assay performed according to the manufacturer’s instructions.

### Quantification of intratubular area

Paraformaldehyde-fixed paraffin-embedded kidney sections were stained with periodic acid-Schiff using standard histological procedures. Intratubular area was quantified from digital images using Fiji. Tubular luminal regions were identified using standard thresholding and segmentation workflows, and intratubular area was calculated for each image as the area enclosed within tubular structures.

### Statistical analyses

All statistical analyses were performed in R^55^. Unless otherwise stated, comparisons between two independent groups were performed using Welch’s *t*-test, while paired *t*-tests were used for repeated measurements from the same animals. Comparisons involving multiple groups or experimental factors were analysed using analysis of variance or linear mixed-effects models, as specified in the corresponding figure legends. Correlations were assessed using Pearson’s correlation coefficient.

For high-dimensional datasets, multiple-testing correction was applied using the Benjamini-Hochberg procedure^56^ unless otherwise specified. Adjusted p values < 0.05 were considered statistically significant.

### Gene and metabolite set analyses

Gene set variation analysis (GSVA) and single-sample directional gene set analysis (ssdGSA) were performed using the GSVA^57^ and ssdGSA^58^ R packages, respectively. ssdGSA was used to perform signed GSVA when a gene set contained a mix of genes with both positive and negative associations to an outcome, enabling clearer insight into the direction of regulation compared with conventional GSVA.

Metabolite set analyses were conducted using the same approaches, with metabolite sets curated as described in the main text.

### Gene set enrichment

Gene set enrichment and over representation analyses were performed using the clusterProfiler package^59^. In over representation tests, the tissue proteome was used as the background to correct for any enrichment relating to the intrinsic function of each tissue.

### Human genetic evidence

Human genetic evidence was acquired and analysed using *Synteny*^11^. Briefly, human genetic evidence scores (HuGE) are calculated by combining the results of many genome-wide association studies to generate a single index for the strength of genetic evidence linking a particular gene to any recorded human phenotype. *Synteny* was used to acquire HuGE scores for genes of interest and assess human phenotype enrichment (HPE). HPE is a statistical approach to determine whether a gene set is more associated with a human phenotype than expected by chance, akin to gene set enrichment.

### Cell-type and functional deconvolution of kidney proteomics

Single-cell RNA-sequencing data from the mouse DKD 1M atlas of Wu et al.^17^ were used as a reference for cell-type and functional deconvolution of bulk kidney proteomics. Cell-type-specific expression signatures were derived from annotated renal cell populations and matched to proteins quantified in the kidney proteome. Relative cell-type proportions were estimated using reference-based decomposition with the Bisque package in R^60^. To assess cell-type-specific functional remodelling, genes differentially expressed between healthy and diabetic kidney disease (DKD) cells were identified separately within each cell type. DKD response signatures were restricted to proteins detected in the kidney proteome and projected onto individual samples using single-sample directional gene set analysis (ssdGSA)^58^. Directional scores incorporated both the magnitude and direction of DKD-associated expression changes, such that higher scores indicated greater similarity to the DKD-associated state for each cell type.

## Data Availability

Proteomics data have been deposited to the ProteomeXchange Consortium via the PRIDE^61^ partner repository. The project accession will be released upon publication.

## Acknowledgements

Thanks go to the technical staff at: Australian BioResources and Laboratory Animal Services at the Charles Perkins Centre of The University of Sydney for providing and maintaining the mice used in this study; the Sydney Imaging core facility at the Charles Perkins Centre for providing assistance in performing EchoMRI; Charles Perkins Centre Histology facility for providing equipment required for microscopic analyses; and Sydney Mass Spectrometry for providing access to liquid chromatography coupled mass spectrometers.

We also thank Dr Óscar Millet and Dr Rubén Gil-Redondo from the Center for Cooperative Research in Biosciences for providing access to the OSARTEN human urinary metabolomics dataset^10^.

## Author contributions

Conceptualisation, HBC, DEJ; Methodology, HBC, ADV, OKF, KCC, SA, KI, TS, LG, FT; Formal Analysis, HBC, ADV, OKF, SA, KI, LG; Investigation, HBC, ADV, OKF, KCC, SWCM, SM, JS, SA, KI, LG; Data curation, HBC, ADV, SA, KI, LG; Visualisation, HBC; Writing – Original Draft, HBC; Writing – Review & Editing, all authors; Supervision, DEJ; Funding Acquisition, DEJ.

## Extended Data Figures

**Extended Data Figure 1.**
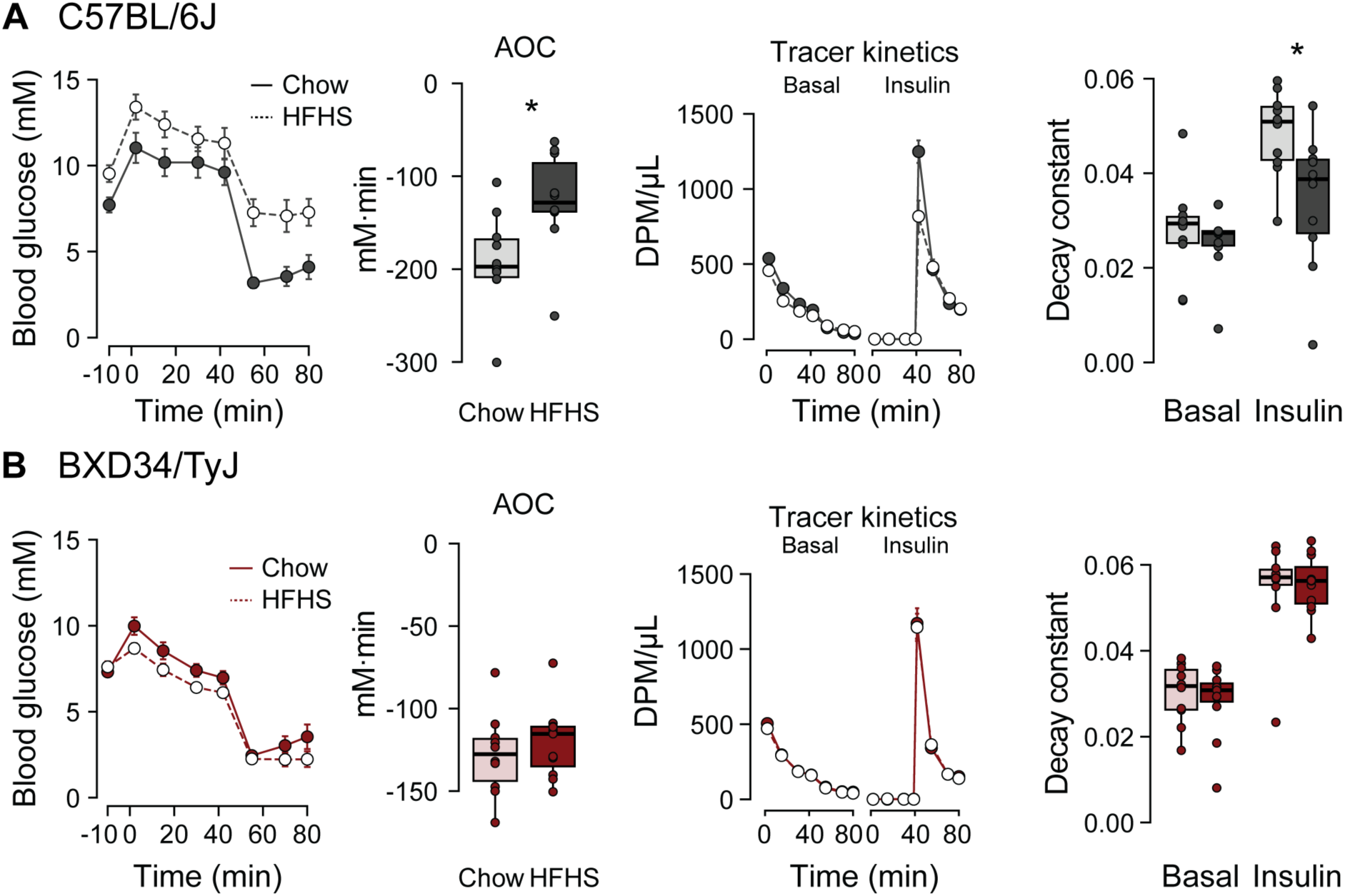
Physiological time course data from the Dual Tracer Test. Supplementary data for **(A)** C57BL/6J and **(B)** BXD34 strains. Left to right: blood glucose concentrations during DTT, insulin administered retro-orbitally at 40 min; area of the curve (AOC) of blood glucose response to insulin; disappearance of radiolabelled 2DG tracers from whole blood; and decay constants for the rate of 2DG disappearance. N = 10/strain/diet. Error bars = SEM. Statistical significance of p < 0.05 is shown as *.

**Extended Data Figure 2.**
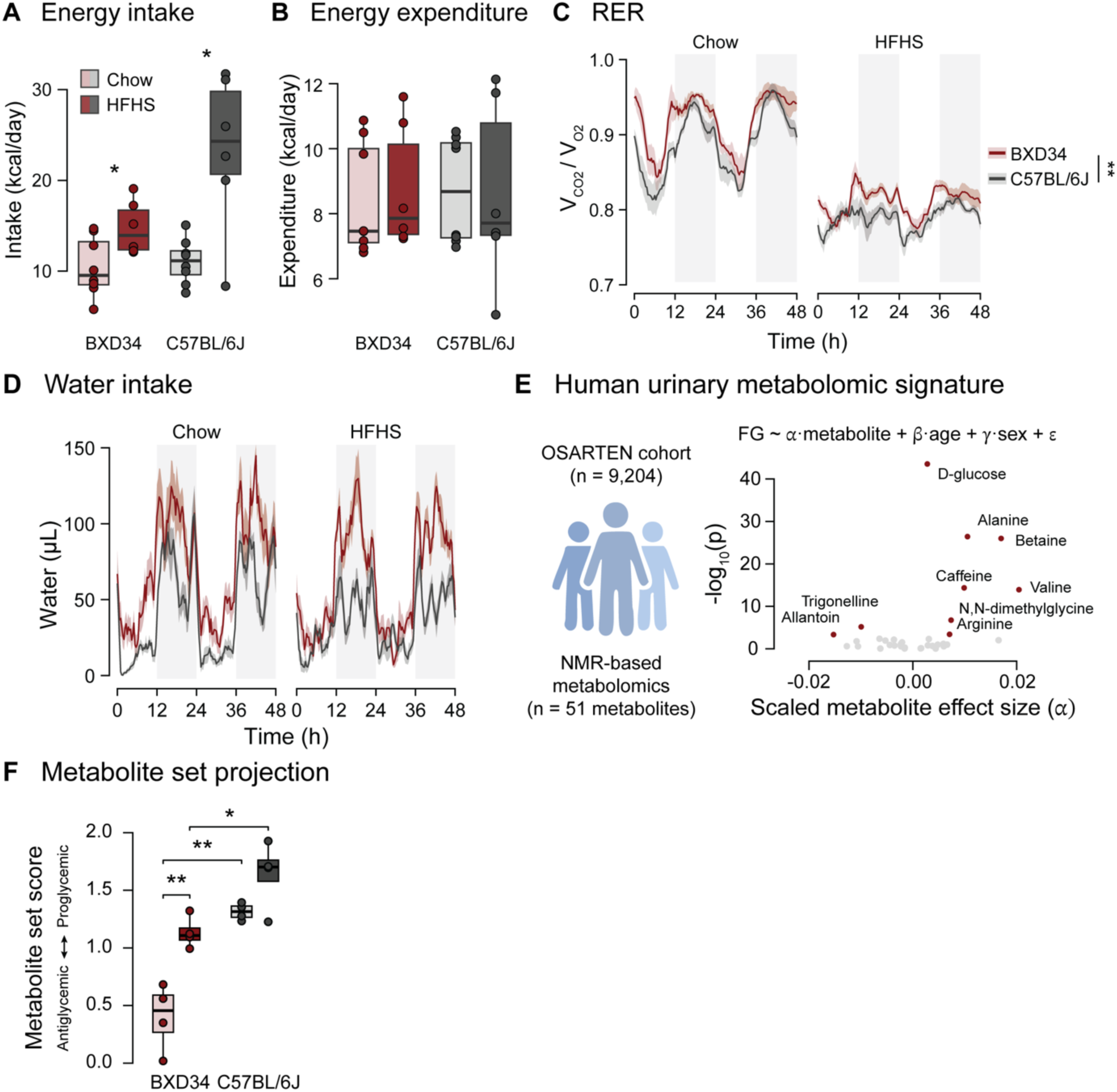
Metabolic cage phenotyping and cross-species urinary metabolite associations. **(A)** Energy intake calculated by multiplying food intake (in grams) by the energy density of the diet (kcal/g). **(B)** Total daily energy expenditure. **(C-D)** Time course measurements of **(C)** respiratory exchange ratio and **(D)** real-time water consumption, presented as a rolling average of measurements ± 30 min. Ribbons represent 95% confidence intervals. Vertical shaded bars represent the dark cycle (1800 hrs – 0600 hrs). N = 8/strain/diet. **(E)** Schematic describing the OSARTEN cohort (left), and results of linear modelling to identify a urinary metabolite signature for fasting blood glucose (FG; right). Significantly associated metabolites are indicated. **(F)** Results of single-sample directional gene set analysis (ssdGSA) using the metabolic signature shown in (E). Higher scores indicate mouse urine samples with increased similarity to urine from humans with hyperglycaemia. Statistical significance of p < 0.05 is shown as *, p < 0.01 as **, and p < 0.001 as ***.

**Extended Data Figure 3.**
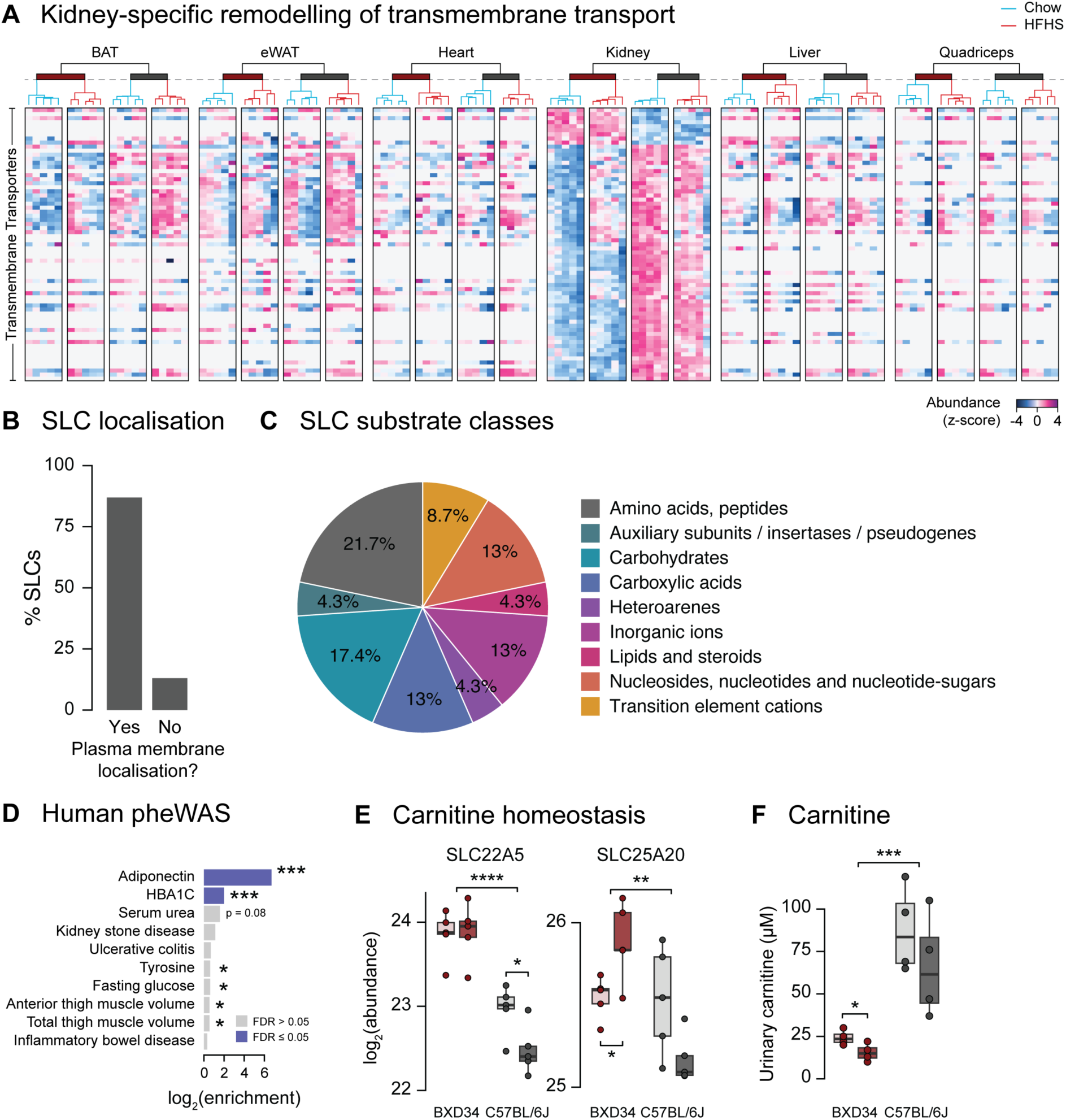
Renal transporter remodelling spans diverse substrates and is linked to human metabolic traits. **(A)** Heatmap of z-scored kidney protein abundances for proteins contained within the Gene Ontology ‘Transmembrane Transport’ gene set that are significantly affected by strain in the kidney. Heatmaps are divided by tissue, strain (indicated by the coloured horizontal bar in the dendrogram), and diet (indicated by the colour of the dendrogram). **(B)** Percentage of SLC proteins downregulated in BXD34 mice with plasma membrane localisation, calculated using the RESOLUTE database^62^. **(C)** Distribution of substrate classes transported by downregulated SLC proteins, calculated using the RESOLUTE database^62^. **(D)** Results of human phenotype enrichment using *Synteny*^11^ to identify human phenotypes with genetic associations to differentially regulated transporters identified in BXD34. Asterisks indicate nominal statistical significance. **(E)** Renal protein abundance of SLC22A5 and SLC25A20, involved in renal carnitine homeostasis. **(F)** Urinary carnitine concentrations in chow- and HFHS-fed C57BL/6J and BXD34 mice. N = 4/strain/diet. Statistical significance of p < 0.05 is shown as *, p < 0.01 as **, and p < 0.001 as ***, and p < 0.0001 as ****.

**Extended Data Figure 4.**
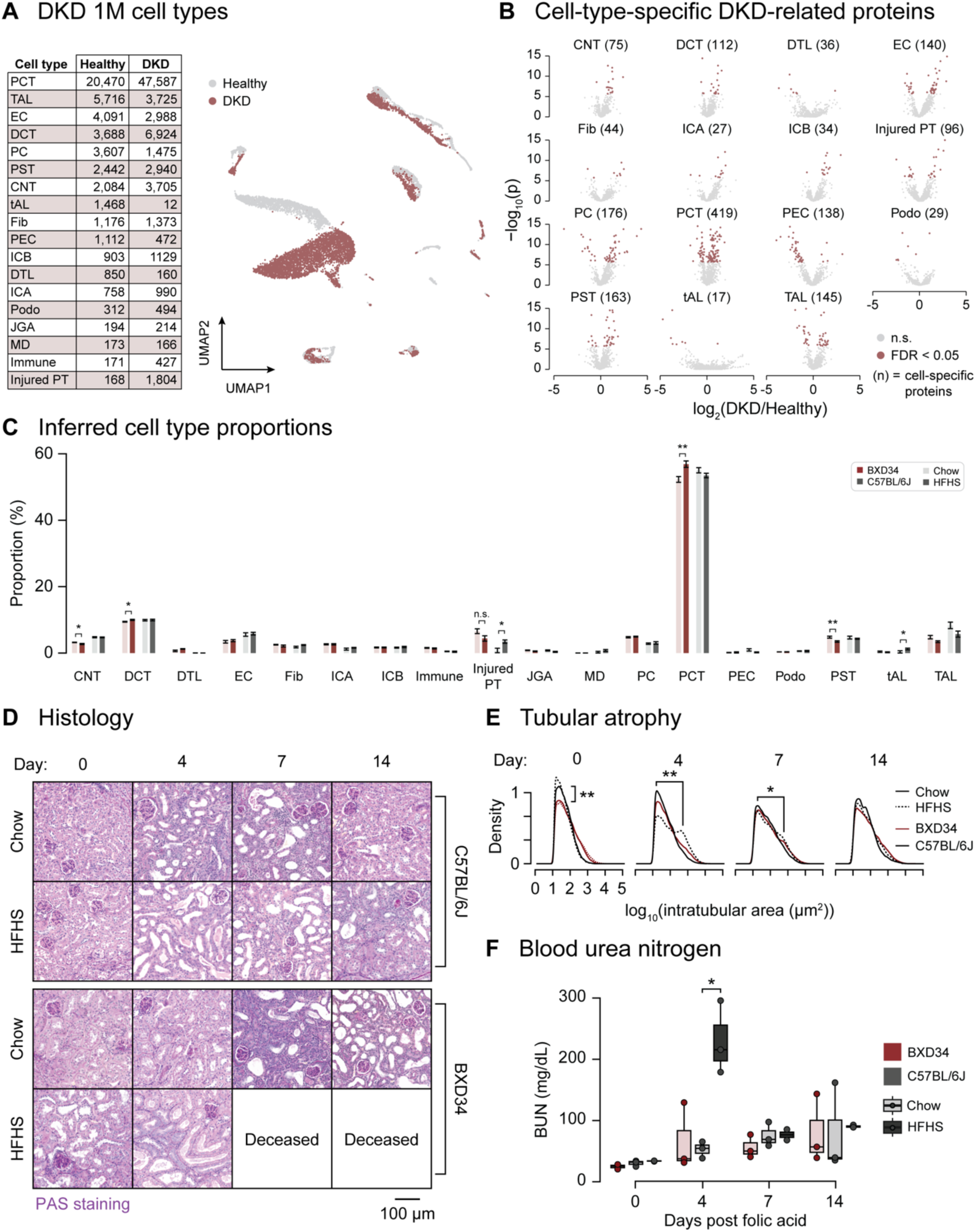
Validation of cell-type deconvolution and renal injury phenotypes. **(A)** Composition of the DKD 1M single-cell RNA-sequencing dataset^17^. Table indicating the number of healthy and DKD cells assigned to each renal cell type (left), and a UMAP demonstrating the distribution of cells derived from healthy and DKD kidneys (right). **(B)** Cell-type-specific DKD-associated proteins used to derive functional signatures. Volcano plots show differential expression between DKD and healthy cells within each renal cell type. Proteins passing FDR < 0.05 are highlighted. Numbers in parentheses indicate the number of cell-type-specific proteins contributing to each signature. Cell types with fewer than 10 unique proteins were excluded from analysis. Cell types: TAL = thick ascending limb; tAL = thin ascending limb; PST = proximal straight tubule; Podo = podocyte; PEC = parietal epithelial cell; PCT = proximal convoluted tubule; PC = principal cell; JGA = juxtaglomerular apparatus; ICB = intercalated B; ICA = intercalated A; Fib = fibroblast; EC = endothelial cell; DTL = descending thin limb; DCT = distal convoluted tubule; CNT = connecting tubule. **(C)** Inferred renal cell-type proportions in chow- and HFHS-fed C57BL/6J and BXD34 mice. Error bars represent SEM. **(D)** Representative PAS-stained kidney sections from chow- and HFHS-fed C57BL/6J and BXD34 mice collected at indicated time points after folic acid administration. No HFHS-fed BXD34 mice survived to days 7 or 14. **(E)** Distribution of intratubular area at indicated time points after folic acid administration, used as an index of tubular atrophy (N = 4 mice/diet/time point). **(F)** Blood urea nitrogen concentrations measured at indicated time points after folic acid administration (N = 3-4/diet/time point). Statistical significance of p < 0.05 is shown as *, and p < 0.01 as **; n.s. = not significant.

